# Anatomical Research on the Anterior Cruciate Ligament of the Knee

**DOI:** 10.1101/2025.11.23.690033

**Authors:** Dongliang Zhang, Zhongbao Han, Yinxiu Chi

## Abstract

**Objective:** The Ribbon Theory challenges the traditional perspective that the anterior cruciate ligament (ACL) consists of a two-bundle structure, specifically the anterior inner bundle and the posterior outer bundle. Consequently, there is a pressing need to conduct further anatomical studies of the ACL to furnish a more precise anatomical foundation for ACL reconstruction procedures.

**Materials and Methods:** A total of 30 fresh knee specimens, sourced from donor cadavers utilized for educational purposes, were dissected. The morphology of the ACL was observed, and its length, insertion position, and surface area were systematically measured.In males, the mean length of the right ACL was 34.52 mm, while the left ACL measured an average of 34.87 mm. In females, the mean length of the right ACL was 30.8 mm, and the left ACL averaged 30.53 mm. The average femoral footprint area on the right side was 143 mm^2^ ± 16 for males and 124 mm^2^ ± 21 for females, whereas on the left side, it was 145 mm^2^ ± 22 for males and 123 mm^2^ ± 17 for females. In conclusion, the majority of ACLs are characterized by a single flat band extending from the femoral origin to the midpoint, with no distinct separation observed at the midpoint. This discovery has significantly influenced the techniques used in ACL reconstruction, advancing the development of the anatomical single-bundle ACL reconstruction method.

## 1 Introduction

The ACL is a critical structure for maintaining knee joint stability, and injuries to the ACL are frequently encountered in clinical practice[1, 2]. ACL surgery involves the reconstruction or replacement of the ligament, with the primary objective being the accurate restoration of the patient’s anatomical structure[3-5]. ACL reconstruction surgery is instrumental in reinstating a painless range of motion, stability, and functionality to the knee following an ACL injury[6, 7]. Conventional research posits that the ACL comprises two distinct bundles: the anteromedial (AM) bundle and the posterolateral (PL) bundle[8]. Historically, ACL reconstruction techniques have predominantly adhered to the two-bundle theory. However, in recent years, the advent of the Ribbon Theory has prompted a reevaluation of the anatomical understanding of the ACL[9-11]. In recent years, the introduction of Ribbon theory has prompted a reevaluation of the anatomical understanding of the ACL[12, 13]. This study seeks to elucidate the precise anatomical characteristics of the ACL by conducting anatomical observations on fresh knee specimens. The findings aim to contribute foundational anatomical insights that could enhance ACL reconstruction techniques. Furthermore, the outcomes of this research are expected to facilitate a more individualized determination of the normal morphology and positioning of the femoral tunnel.

## 2 Materials and methods

### 2.1 Ethics and Samples

The recruitment period of this study began on January 1, 2021 and ended on December 31, 2024. Participants have provided informed consent orally.The study received approval from the Institutional Review Committee and Ethics Committee of the “Research Platform of Jiangsu Medical University.” This anatomical, observational, and comparative investigation involved 15 unmatched, formalin-fixed human cadaveric knees, sourced from donors utilized for educational purposes at medical colleges in China. All specimens were from individuals who were over 18 years of age at the time of death. Detailed records of each cadaver’s sex, weight, and height were maintained. Additionally, the cause of death and pertinent clinical data for each cadaver were documented, neither the cause of death nor the medical history of the cadavers included in the study influenced the methodology or outcomes. During the anatomical examination, we meticulously observed the morphology and insertion sites of the ACL and employed precise measuring instruments to determine its length. Subsequently, the images were transferred to a personal computer, where the footprint area was quantified using ImageJ software. All measurement data underwent statistical analysis to ensure the accuracy and reliability of the findings.

### 2.2 Cadaver perfusion method

The composition of the traditional perfusion solution is as follows: 37% formaldehyde solution at 10% (v/v), 95% ethanol at 15% (v/v), glycerol at 2% (v/v) for moisturizing and preventing tissue shrinkage, sodium chloride at 0.9% (w/v) to maintain isotonic conditions, and distilled water to complete the solution to 100%. The cadaver is positioned supine with a slight abduction of the hip joint. A 5 cm longitudinal incision is made 1 cm inferior to the inguinal ligament to expose the common femoral artery (CFA) through blunt dissection. The proximal end of the artery is secured with a 2-0 suture, while the distal end is temporarily ligated. A 16G venous catheter or an 8F arterial sheath is inserted into the CFA in the direction of the heart for a distance of 3–4 cm and is secured upon confirmation of blood return. Perfusion is conducted either manually using a 50 mL plastic syringe or via gravity drip from a height of 80 cm, equivalent to 60 mmHg. Initially, 20 mL of heparin-saline solution (5 IU/mL) is rapidly injected to clear any blood clots, followed by the administration of the traditional perfusion solution. The total volume of the perfusion solution is calculated as 2% of the body weight, approximately 150–200 mL for a 70 kg adult. The catheter is removed once the toenail bed becomes pale and subcutaneous superficial veins are visible. The proximal end of the artery is then double-ligated using silk sutures.

### 2.3 Statistical Analysis

The statistical analysis was conducted using the SPSS software package. Data are presented as mean ± standard deviation (SD). Intergroup differences were evaluated using Student’s t-test, with p-values less than 0.05 considered statistically significant.

## 3. Results

### 3.1 Construction of Perfusion System

A closed extracorporeal constant-flow perfusion model (Figure 1A) was employed in this study. The cadaver was positioned supine, and a longitudinal incision measuring 8 cm was made at the midpoint of the inguinal region to expose the common femoral artery (CFA) (Figure 1B). The pump head speed was adjusted to 0.6–1.2 mL min^−1^, representing 10% of the physiological flow rate of the CFA (60–80 mL min^−1^), in order to minimize mechanical damage to the vascular endothelium as much as possible.

**Figure 1.**
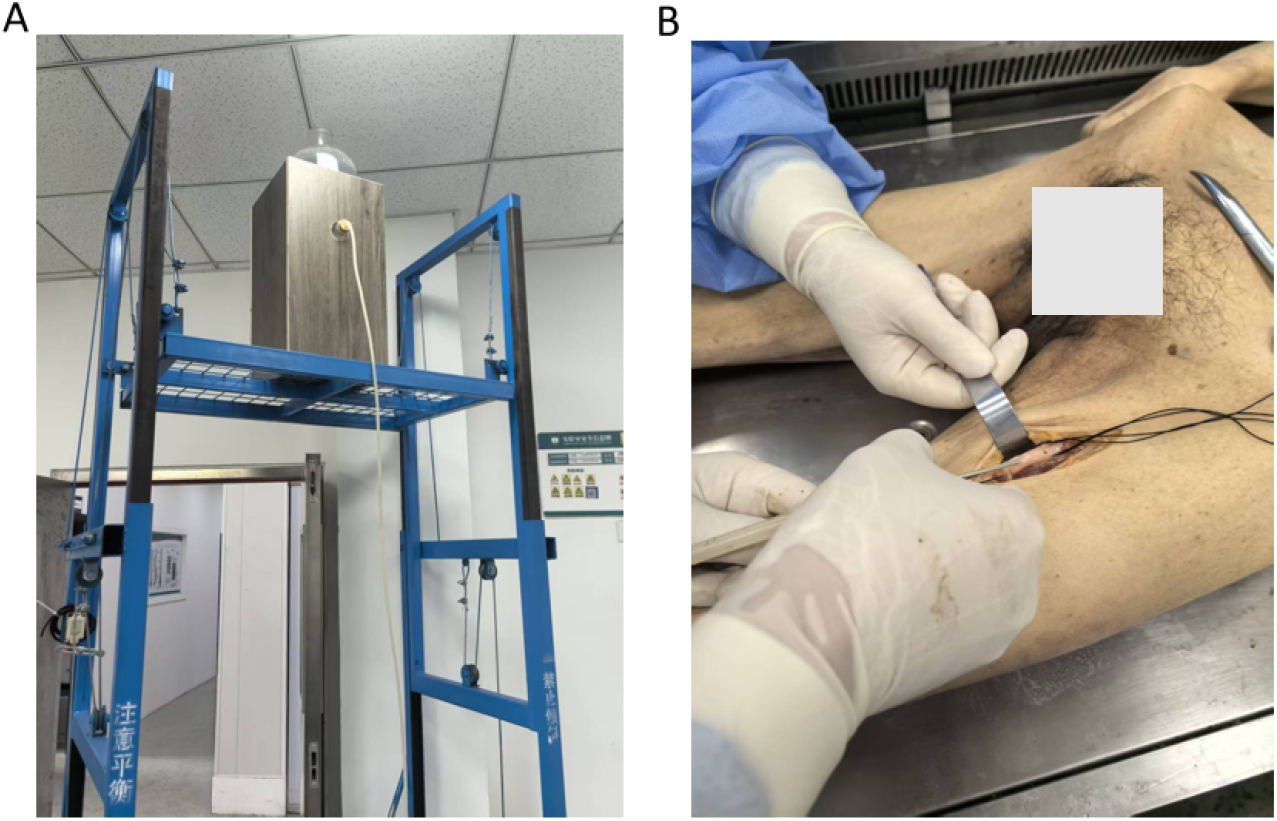
Cadaver perfusion procedure. (A) Self-made electric infusion equipment (B)Perform puncture on the exposed femoral artery.

**Figure 2.**
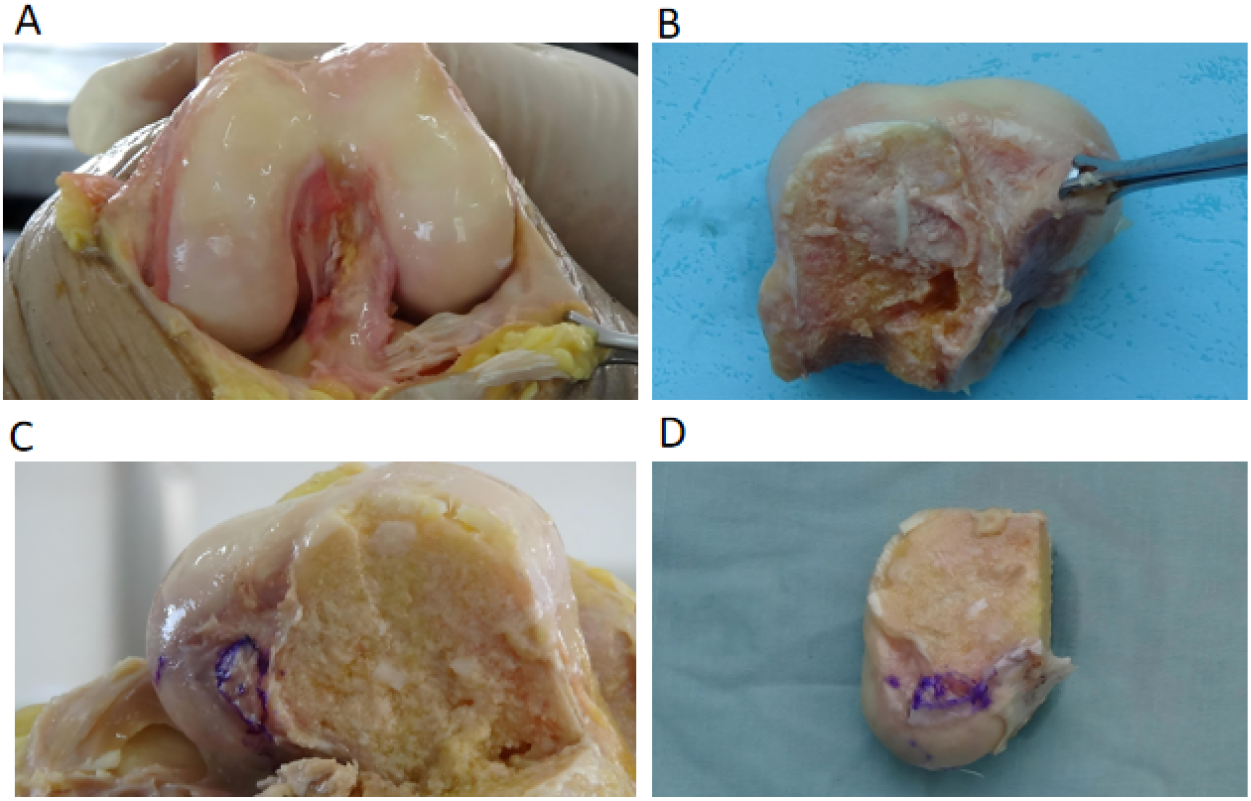
CIllustrates the anatomical morphology of the ACL in a cadaveric knee: (A,B) a single strand of severed ACL, (C) the femoral attachment of a single strand of ACL, and (D) the partial attachment of a double strand ACL.

### 3.2 Observation of ACL Morphology

The study results indicated that 24 out of 30 knees exhibited a single flat band extending from the femoral origin to the midsection, without any discernible separation in the middle region. The femoral footprint was observed to be oval, which contrasts significantly with the traditional two-bundle theory (Fig. 1A, B). Only six ACLs displayed a banded structure divided into two bundles, although the demarcation between these bundles was not distinct (refer to Table 1). Furthermore, only two distinct insertions were identified at the femoral attachment site. Observation of ACL Femoral Footprint: The single ACL femoral footprint was characterized as oval (Fig.1C);The femoral footprint of the double ACL was delineated. (Fig1D).

**Table 1.** Distribution of the sample by gender of the knees. presents the p-values obtained from comparing the embedded shape relative to gender.

| Table 1 Distribution of the sample by gender of the knees |  |  |  |
| --- | --- | --- | --- |
| Category | semicircular | Strip shape | Female vs. male<br>(*p* value) |
| Male left<br>(n=9) | 7 | 2 | n.s. |
| Female left<br>(n=6) | 5 | 1 |  |
| Male right<br>(n=9) | 8 | 1 | n.s. |
| Female right<br>(n=6) | 4 | 2 |  |

ACL length measurements indicate that the average length of the left ACL in males is 34.87 mm, while the right ACL averages 34.52 mm. In females, the average length of the left ACL is 30.53 mm, and the right ACL averages 30.8 mm.The lengths of the ACL varied between sexes and sides, although these differences did not reach statistical significance (Table 2). Regarding the measurement of the ACL femoral footprint area, the average area on the left side was 145 mm^2^ ± 22 for males, compared to 143 mm^2^ ± 16 on the right side. For females, the average area was 123 mm^2^ ± 17 on the left and 124 mm^2^ ± 21 on the right. Notably, there were statistically significant differences in the ACL femoral footprint area between genders and sides (Table 2).

**Table 2.** Comparison between the average values obtained in each gender. Values of *p* obtained by comparing the length and area relative to gender

| Table 2 Comparison between the average values obtained in each gender |  |  |  |
| --- | --- | --- | --- |
| Category | Length (mm) | Area (mm<sup>2</sup>) | Female vs. male (p value) |
| Male left (n=9) | 34.87 | $145 \pm 22$ | Length: n.s.<br>Area: <0.05 |
| Female left (n=6) | 30.53 | $123 \pm 17$ | |
| Male right (n=9) | 34.52 | $143 \pm 16$ | Length: n.s.<br>Area: <0.05 |
| Female right (n=6) | 30.8 | $124 \pm 21$ | |

## 4 Discussion

Scientific evidence suggests that double bundle anterior cruciate ligament reconstruction (DBACL) more accurately replicates the native anatomy of the ACL compared to single bundle reconstruction[14]. Nonetheless, the extent to which DBACL results in superior functional outcomes remains a topic of ongoing debate[15, 16]. Despite this, anatomic single bundle techniques have traditionally been regarded as the gold standard for ACL reconstruction[17, 18].

ACL is integral to the stabilization of the knee joint, particularly during activities that involve stopping, turning, and changing direction. The debate regarding whether the ACL comprises two bundles or a single bundle is primarily pertinent to the selection of grafts in ACL reconstruction surgery[19]. During such procedures, surgeons must decide between employing a “double-bundle” or a “single-bundle” graft technique. Double-bundle reconstruction involves the use of two grafts inserted through two separate bone tunnels[20], whereas single-bundle reconstruction utilizes a single graft[21]. The distinction between “double bundle” and “single bundle” grafts extends beyond the mere difference in the number of strands. A single bundle graft typically comprises a four-strand configuration, created from two tendons folded in half[22], whereas a double bundle graft results in an eight-strand structure[23]. From a mechanical strength perspective, the normal ACL exhibits a tensile strength of approximately 2000 newtons, whereas an individual tendon typically demonstrates a strength of around 1000 newtons. Notably, the mechanical strength of the commonly employed single bundle graft, which consists of four strands derived from the femoral tendons, surpasses 4000 newtons, Theoretically, the mechanical strength of a single-bundle graft is adequate to meet clinical requirements[13]. Although double-bundle reconstruction offers superior mechanical strength and enhanced anti-rotational stability, long-term clinical follow-up studies have demonstrated that patients undergoing double-bundle reconstruction exhibit knee function and re-failure rates comparable to those undergoing single-bundle reconstruction[9]. Furthermore, double-bundle reconstruction is often associated with increased surgical trauma, higher financial costs, and challenges in surgical revision following re-injury. Consequently, not all patients are ideal candidates for double-beam reconstruction. Currently, over 80% of clinicians predominantly opt for single-beam reconstruction, achieving satisfactory clinical outcomes[22].

The findings of this study indicate that anterior cruciate ligaments (ACLs) are primarily single-bundle flat banded structures, as opposed to the conventional double-bundle configurations. This discovery holds significant implications for ACL reconstruction methodologies. The traditional double-beam reconstruction technique may be overly complex, without offering superior efficacy compared to single-beam reconstruction. Consequently, anatomic single-bundle anterior cruciate ligament reconstruction (ASB-ACLR) has gained prominence due to its relative ease of execution and its superior ability to restore knee joint stability. Furthermore, variations in the length of the ACL and the femoral footprint have been observed between genders and between the left and right sides. These anatomical differences may influence the techniques employed in ACL reconstruction, underscoring the necessity of considering patient-specific anatomical variations when devising surgical plans.

## 5 Conclusion

Through anatomical examination of 30 fresh knee specimens, this study elucidated that the true morphology of the ACL is that of a single bundle of flat band structure. This discovery significantly influences the reconstruction of the ACL and advances the development of anatomical single-bundle ASB-ACLR echnology. Future research should further investigate the relationship between the anatomical characteristics of the ACL and reconstruction techniques to offer more precise and effective solutions for treating ACL injuries.

## Acknowledgments

We extend our gratitude to all donors and their families for providing fresh knee specimens for this study. Additionally, I express my appreciation to the research team members for their diligent efforts and selfless dedication. Through anatomical examination of 30 fresh knee specimens, this study substantiated the validity of the Ribbon theory, which posits that the ACL is a single flat band structure extending from the femoral origin to the midsection. This finding has significantly influenced ACL reconstruction techniques and has advanced the development of anatomical single-bundle ACL reconstruction methods. Future research should further investigate the clinical efficacy of single-bundle anterior cruciate ligament reconstruction techniques to offer more effective treatment options for patients with ACL injuries.

## Author contributions

Design of the project was contributed by Dongliang Zhang, Data collection was contributed by Zhongbao Han; Writing and editing were contributed by Yinxiu Chi.

## Data availability

The datasets used and/or analysed during the current study available from the corresponding author on reasonable request.

